# High sensitivity dia-PASEF proteomics with DIA-NN and FragPipe

**DOI:** 10.1101/2021.03.08.434385

**Authors:** Vadim Demichev, Fengchao Yu, Guo Ci Teo, Lukasz Szyrwiel, George A. Rosenberger, Jens Decker, Stephanie Kaspar-Schoenefeld, Kathryn S. Lilley, Michael Mülleder, Alexey I. Nesvizhskii, Markus Ralser

## Abstract

The dia-PASEF technology exploits ion mobility separation for high-sensitivity analysis of complex proteomes. Here, we demonstrate neural network-based processing of the ion mobility data, which we implement in the DIA-NN software suite. Using spectral libraries generated with the MSFragger-based FragPipe computational platform, the DIA-NN analysis of dia-PASEF raw data increases the proteomic depth by up to 69% compared to the originally published dia-PASEF workflow. For example, we quantify over 5200 proteins from 10ng of HeLa peptides separated with a 95-minute nanoflow gradient, and over 5000 proteins from 200ng using a 4.8-minute separation with an Evosep One system. In complex samples, featuring a mix of human and yeast lysates, the workflow detects over 11700 proteins in single runs acquired with a 100-minute nanoflow gradient, while demonstrating quantitative precision. Hence, the combination of FragPipe and DIA-NN provides a simple-to-use software platform for dia-PASEF data analysis, yielding significant gains in high-sensitivity proteomics.

## Introduction

Data-independent acquisition (DIA) methods in proteomics (Gillet et al., 06/2012; Venable et al., 2004) have been actively developed in the past few years, leading to increased depth, higher data consistency and accurate quantification (Ludwig et al., 2018). DIA methods have become attractive in classic proteomics that relies on conventional nano-flow chromatography, where they increase depth in ‘single-shot’ experiments. For example, a recent study demonstrated quantification of more than 10000 proteins in human cell lysates (Muntel et al., 2019). Further, significant progress in the analysis of DIA data has allowed it to move to faster gradients and higher flow rates. These approaches benefit from higher peak capacities, increased column lifetime, improved chromatographic stability and higher throughput, but at the expense of a higher sample dilution (Bian et al., 2020; Messner et al., 2020b; Vowinckel et al., 2018). These developments now enable applications where throughput, consistent quantification and low batch effects are essential. These applications include, among others, exploratory drug screens and clinical studies with high participant numbers. For example, we have recently demonstrated SWATH-MS experiments that use 5-minute 800μL/min high flow chromatography and can maintain high quantification accuracy over thousands of samples (Demichev et al., 2020; Messner et al., 2020b). More recently, the introduction of Scanning SWATH has facilitated sub-minute proteomics, and allowed for the precise quantification of about 2700 protein groups using 60-second chromatographic gradients (Messner et al., 2020a).

Both conventional and ultra-fast proteomics can further gain depth and quantitative accuracy with additional separation of the convoluted precursor or fragment ion space. A recent acquisition method, dia-PASEF, utilizes the Trapped Ion Mobility Separation (TIMS) device within the timsTOF Pro mass spectrometer (Bruker Daltonics) (Meier et al., 2020). In dia-PASEF, the ion mobility dimension allows to distinguish signals from peptides that would otherwise be co-fragmented, thus producing cleaner spectra. Most importantly, dia-PASEF yields 2-to 5-times improvement in sensitivity, depending on the acquisition scheme, by ‘stacking’ precursor ion isolation windows in the ion mobility dimension and thus increasing the effective duty cycle.

These features render dia-PASEF complementary to the recent Scanning SWATH acquisition technique (Messner et al., 2020a). The TIMS ion mobility separation is slower than the fast tandem-MS scans of Scanning SWATH, and dia-PASEF is hence less suited for being used in conjunction with ultra-fast chromatographic gradients generated with analytical flow LC systems (Messner et al., 2020a, 2020b). However, the ion mobility separation and the resulting boost in sensitivity can maximize proteomic depth when only limited sample amounts are available, such as in single cell proteomics (Brunner et al., 2020) or in the quantitative analyses of post-translational modifications (Bekker-Jensen et al., 2020; Hansen et al., 2021; Steger et al., 2020).

Here we present a workflow for dia-PASEF data processing that provides a significant improvement in the depth of proteome coverage and quantitative precision. We have incorporated a software solution for the processing of TIMS ion mobility dimension through the use of deep neural networks in DIA-NN, an automated software suite that also simplifies and accelerates DIA data analysis (Demichev et al., 2020). Moreover, we have optimized the generation of spectral libraries from offline-fractionated PASEF (DDA) data using FragPipe computational platform. FragPipe provides fast, sensitive, and fully automated solution for the analysis of DDA data, from peptide identification with MSFragger search engine (Kong et al., 2017; Yu et al., 2020a), to peptide validation, protein inference and false discovery rate (FDR) based filtering using Philosopher (da Veiga Leprevost et al., 2020), to effective nonlinear retention time and ion mobility alignment between different DDA runs and generation of the final spectral library with EasyPQP. We show that the analysis of dia-PASEF data with this extended DIA-NN version and FragPipe-generated libraries increases the proteomic depth of existing raw data by up to 69%, while simultaneously increasing data consistency as well as quantification accuracy and precision.

## Results

We have designed and incorporated a TIMS module in DIA-NN (Demichev et al., 2020), which allows to extract ion mobility-separated data, characterise the quality of peptide-spectrum matches using the agreement between the expected and observed ion mobility values, and assess these quality scores using neural networks (Figure 1). In contrast to the direct extraction of ‘profile’ ion mobility data as implemented in the Mobi-DIK module for OpenSWATH (Meier et al., 2020), DIA-NN starts the analysis from 2-dimensional peak-picking, wherein a scanning window is used to find local maxima in the 1/K0 x m/z space, where 1/K0 is the inverse ion mobility. Importantly, this algorithm does not produce centroided data, but instead can represent a noisy signal corresponding to a peptide as several closely located peaks, when the real peak’s m/z and 1/K0 values are difficult to identify with high confidence. Among these peaks, different ones might be eventually used during subsequent targeted chromatogram extraction, depending on the reference ion mobility and fragment masses of a particular query precursor ion, thus maximising the sensitivity. Specifically, for each precursor ion and for each of its fragment ions, the most intense peaks are identified within a particular mass threshold (automatically determined or user-defined) and ion mobility threshold (automatically determined based on the alignment between confident identifications and the spectral library). Once whole chromatograms are extracted for a specific precursor and its fragments, candidate peaks, featuring signals from at least two different fragments, are identified and scored. The consistency of ion mobility values between fragment ions is taken into account, and ‘outlier’ fragments get lower scores, even if their elution profiles show high correlation with those of other fragments. This eventually leads to preferential selection of candidate elution peaks with high consistency of ion mobility values across fragments. Further, the deviation of observed fragment ion mobilities from the reference precursor ion mobility value obtained from the spectral library is likewise taken into account. Ultimately, these scores are fed into the ensemble of deep neural networks that DIA-NN uses to assign confidence scores to precursor-spectrum matches (Demichev et al., 2020). Finally, signals with deviating ion mobility values are excluded during the quantification of peptides, thus improving quantitative accuracy (Figure 1).

**Figure 1.**
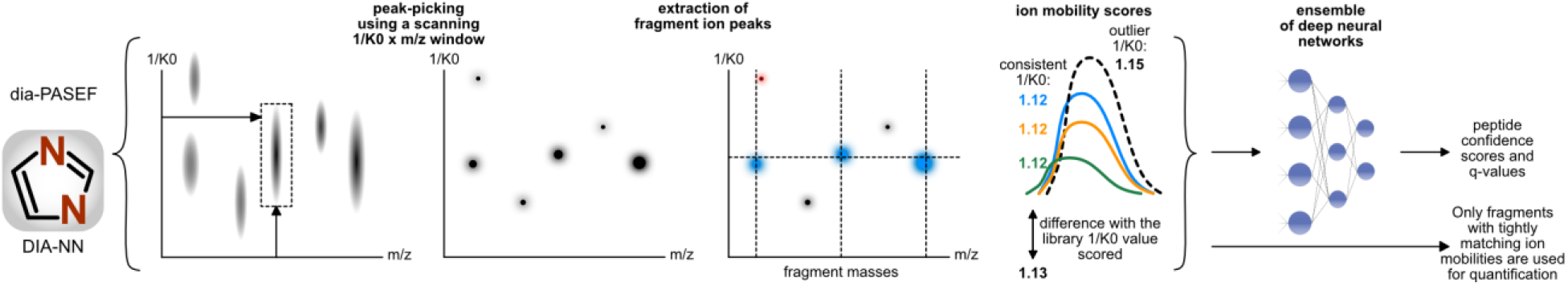
Ion mobility data processing implemented in DIA-NN. The ion mobility module in DIA-NN implements 2D-peak-picking, followed by chromatogram extraction, wherein for each precursor or fragment ion only peaks within certain m/z and ion mobility thresholds from the expected values are used. Inverse ion mobility values (1/K0) are compared between different fragment ions (extracted elution profiles of which are indicated with different colors) as well as to the reference library 1/K0 value, to score putative peptide-spectrum matches. The resulting data is analysed by an ensemble of deep neural networks, to distinguish true and false signals. Signals with deviating ion mobility values are also filtered out to increase quantification accuracy.

Next to the ability to make use of deep neural networks in the analysis of the ion mobility dimension in DIA-NN, we speculated that further grains in dia-PASEF performance can be obtained with an adapted workflow for the generation of specific spectral libraries. We extended our earlier work (Yu et al., 2020a) and optimized a data analysis workflow (see Methods), now available as one of the default workflows in FragPipe, for generating spectral libraries from fractionated PASEF data in the format directly compatible with DIA-NN.

To benchmark the performance of the ion mobility module in DIA-NN in combination with the FragPipe-generated libraries, we reprocessed the dia-PASEF reference dataset (Meier et al., 2020), wherein a HeLa tryptic digest was acquired using different injection amounts (10ng - 100ng on a Thermo Fisher EASY-nLC 1200 nanoLC, 95-minute gradient) and chromatographic gradient lengths (200ng, 4.8-min to 21-min on an Evosep One microflow system) (Figure 2a). In the nanoflow experiments, the FragPipe-generated library built from 24 fractionated DDA PASEF runs and filtered at 1% protein and peptide FDR contained 9991 proteins and 161325 peptides (see Table 1 and Methods). Using this spectral library, we quantified 5227 proteins per run from 10ng of HeLa peptides as well as 7195 proteins from 50ng, out of which 7104 were uniquely identified using proteotypic peptides. This is a gain of 731 proteins compared to the analysis of 100ng injections in the same experiment with the original software workflow (Meier et al., 2020). Furthermore, our computational pipeline performed well with both acquisition schemes used, the 25% duty cycle scheme and the standard scheme (Meier et al., 2020). The 25% duty cycle scheme was more precise quantitatively, suggesting that it is likely preferable for analyses of low sample amounts (Figure 2b).

**Figure 2.**
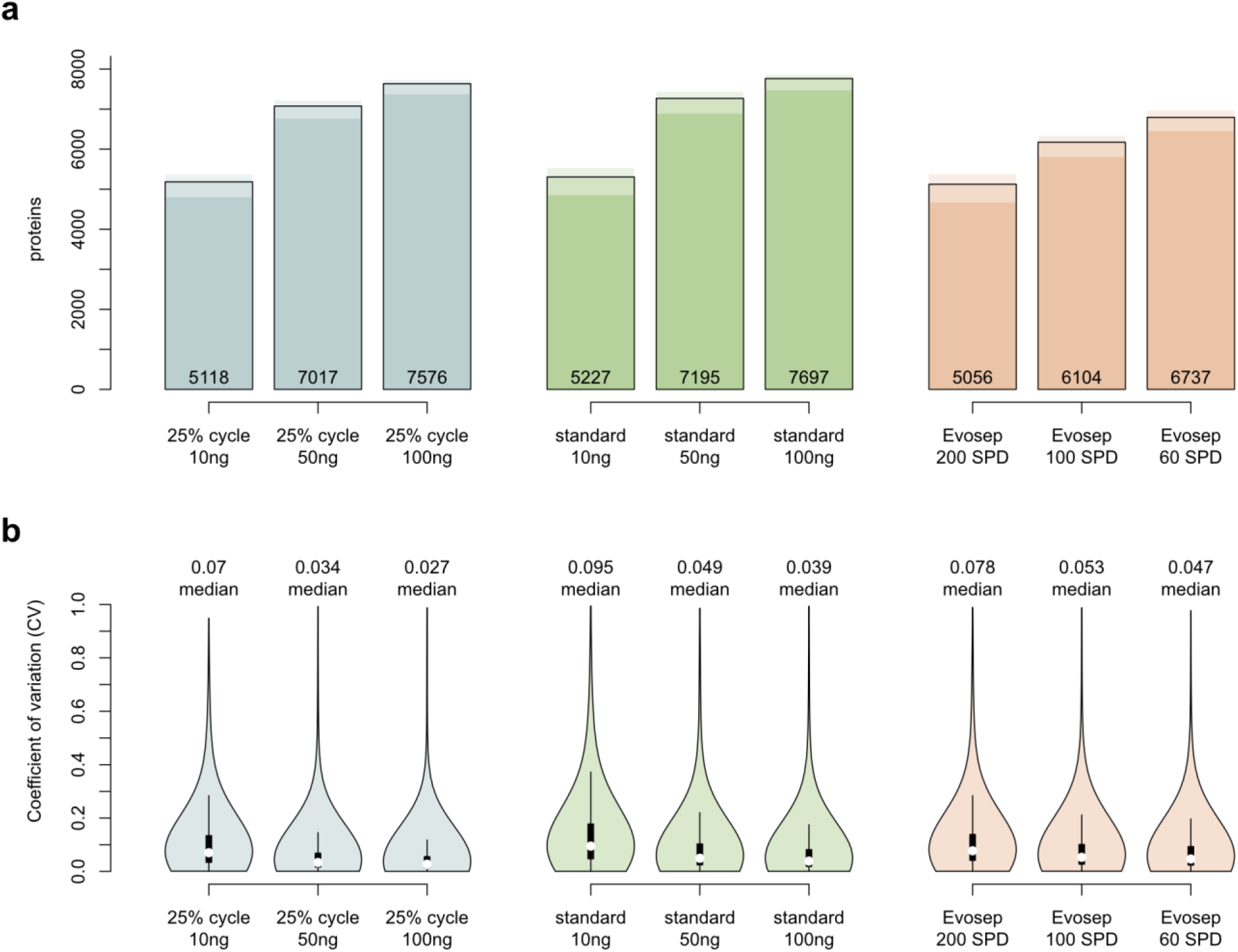
Protein detection and quantification precision. **a** Number of quantified proteins for different injection amounts and instrument settings. Numbers of proteins detected in 1, 2 or all 3 replicates for each dataset (nanoflow 25% duty cycle scheme and standard scheme; Evosep 200, 100, and 60 samples per day (SPD) methods) are shown with different color shades. **b** Coefficients of variation (CV) distributions for the same datasets.

**Table 1.** Spectral libraries.

| Library | Ions | Peptides | Proteins | Genes |
| --- | --- | --- | --- | --- |
| HeLa, nanoflow | 260785 | 161325 | 9991 | 9973 |
| HeLa, nanoflow, semi | 443649 | 306564 | 10177 | 10158 |
| HeLa, Evosep One | 145875 | 98485 | 8201 | 8187 |
| HeLa, Evosep One, semi | 149188 | 102307 | 8248 | 8233 |
| HeLa, two-organism | 361555 | 224597 | 10353 | 10332 |
| HeLa, two-organism, filtered | 360458 | 223907 | 10350 | 10331 |
| Yeast, two-organism | 134148 | 79860 | 5134 | 5132 |
| Yeast, two-organism, filtered | 133351 | 79337 | 5113 | 5113 |

DIA-NN had originally been conceived to maximize the performance of fast proteomic experiments, an area that was previously not a focus of the majority of DIA software developments (Demichev et al., 2020). Thus, we sought to evaluate the performance of DIA-NN with FragPipe-generated libraries on dia-PASEF data with short chromatographic gradients acquired (Meier et al., 2020) using the Evosep One system (Bache et al., 2018). The spectral library built with FragPipe from 24 DDA PASEF runs contained 8201 proteins, 98485 peptides, and 145875 precursor ions (Table 1). Using this library, DIA-NN quantified on average 5056 proteins from 200ng of HeLa digest analysed with a 4.8-minute gradient (200 SPD (samples per day) Evosep One method). Out of these proteins, 4965 were uniquely identified using proteotypic peptides, with 69% improvement over the current state of the art (Meier et al., 2020). The 4.8-minute gradients thus yielded a number of quantified proteins comparable to that reported for 21-minute gradients (60 SPD method) in the original publication (4965 vs 4954).

We then performed a number of additional analyses to confirm the validity of observed gains and to investigate the major reasons for improvement. First, using a two-species benchmark strategy (Muntel et al., 2019), in dia-PASEF data with varying levels of complexity, we confirmed that the gains in proteomic depth are not caused by an inflation of the FDR. Indeed, the analysis revealed that the FDR values reported by DIA-NN are conservative (Supp. Figure S1). We also processed the same dia-PASEF data with DIA-NN using the original spectral libraries (Meier et al., 2020), to quantify the contribution of the FragPipe libraries to the gains in proteomic depth (Supp. Figure S2a).

Despite the increase in the depth of proteome coverage, parameters important for robustness and quantification precision were not compromised, and indeed, did improve in comparison to the original values. Most notably, we obtained high data completeness, which renders the workflow attractive for the application of machine learning methods in the analysis of large sample series, that strongly profit from consistent data. Data completeness ranged from 94% for 200 SPD Evosep One benchmark to 98% for 100ng injections with the 25% duty cycle scheme. Moreover, expressed as Coefficient of Variation (CV), the quantification precision was high - especially for datasets with higher injection amounts (median CV of 2.7% for the nanoflow, 25% cycle, 100ng injection experiment) - and the median CVs were better than 10% across all datasets (Figure 2b). The technical variation in the Evosep One benchmarks here reflects not just the variability introduced by chromatography and mass spectrometer performance, but also that due to peptide purification via solid-phase extraction in Evotips. Nevertheless, a median CV value below 8% was achieved even for the ultra-fast 200 samples per day Evosep One method.

We also sought to gain a better understanding of the degree of gas-phase fragmentation prior to MS/MS, which we previously observed in timsTOF PASEF data (Yu et al., 2020a), and how it may affect the depth of coverage and the quantification accuracy of dia-PASEF. Thus, we constructed a spectral library for HeLa nanoflow data using FragPipe with MSFragger semi-enzymatic search. We observed a very high number of semi-tryptic peptides (Table 1), and a modest increase in the number of identified proteins (10177 vs. 9991 in fully enzymatic search). While the use of this much larger semi-tryptic spectral library in DIA-NN led to identification of 11%, 19% or 24% more precursors on average, for 10ng, 50ng and 100ng injections with 25% duty cycle acquisition scheme, it did not translate into a gain in the number of quantified proteins, while the CV values remained comparable (Suppl. Figure S2), presumably due to overall low intensity of ions generated via gas-phase fragmentation. Of note, we observed a significantly lower rate of gas-phase fragmentation events in recent Evosep One datasets (Table 1). Overall, we conclude that the gas-phase fragmentation had minor impact and accounting for it during data analysis is unlikely to be beneficial.

Finally, we analysed the quantification accuracy using the two-organism benchmark data (Figure 3), wherein a yeast (*Saccharomyces cerevisiae*) lysate was spiked in different concentrations (45ng, sample A, and 15ng, sample B) into a HeLa lysate (200ng) and analysed with a 100-minute nanoLC gradient on a Bruker nanoElute LC system (Meier et al., 2020). The human and yeast spectral libraries built with FragPipe from 24 (HeLa) and 48 (yeast) PASEF runs contained 10353 and 5134 proteins, respectively (Table 1). On average, we report the quantification of 11732 proteins per run for sample A and 11392 for sample B. The identification of yeast proteins in this benchmark is particularly challenging for the low-concentration sample B (15ng yeast lysate only), and thus the numbers of valid A:B ratios reflect the ultimate sensitivity of the workflow. We report 2988 valid A:B ratios for yeast proteins, exceeding the previously reported number of 1394 for the same experiment (Meier et al., 2020). The DIA-NN workflow also produced better overall quantification accuracy, with visibly less grossly incorrect A:B ratios for yeast proteins. The numbers of human protein ratios were moderately higher (8671, +13%), approaching the limit (~10000) of expressed proteins detectable by mass spectrometry in HeLa cells (Muntel et al., 2019). For these, a quantification precision of 3.6% median CV was observed. In total, 11795 proteins were quantified in the whole experiment.

**Figure 3.**
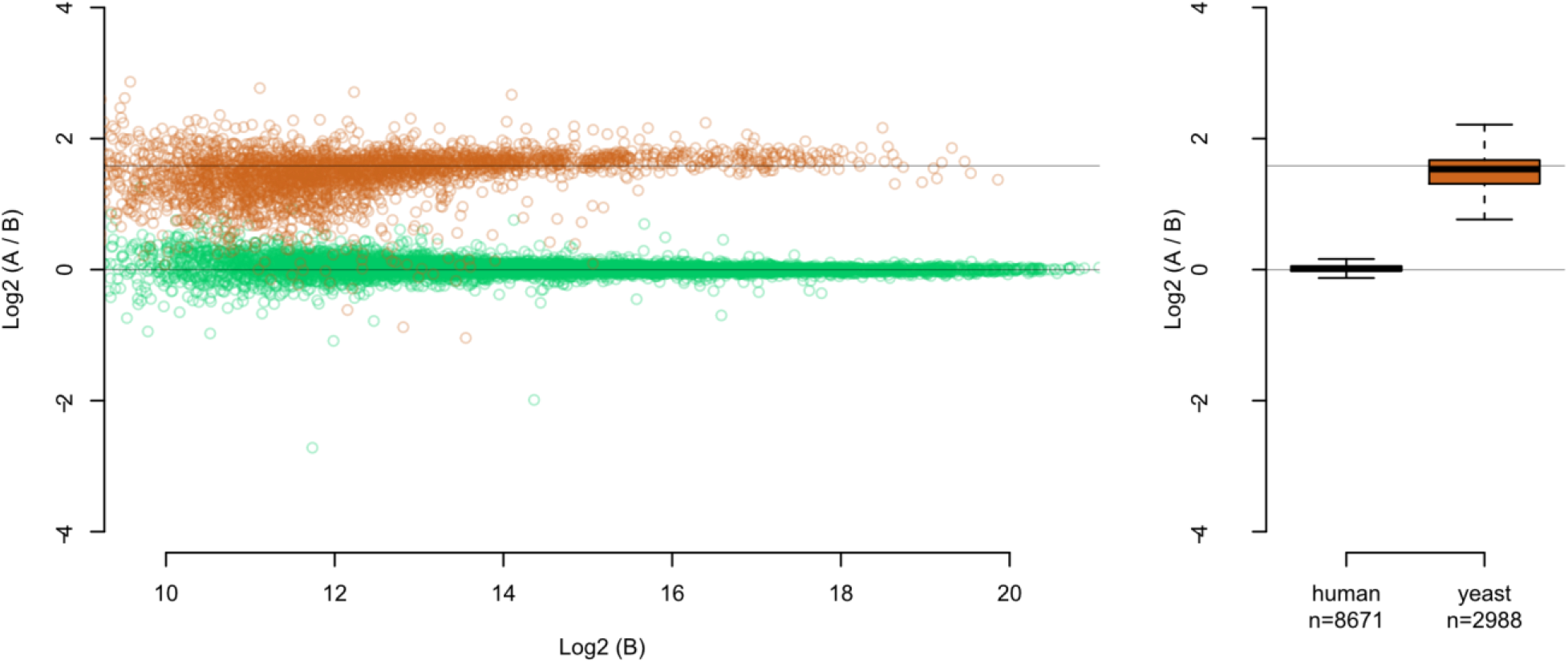
Quantification accuracy. A yeast digest was spiked into a HeLa digest (200ng) in different proportions (A, 45ng and B, 15ng) and analysed in triplicates using a 100-minute nanoLC gradient (Meier et al., 2020). The runs were processed with DIA-NN using a spectral library created with FragPipe. On the boxplot, the boxes correspond to the interquartile range, and the whiskers extend by 1.5x interquartile range. Expected ratios are indicated with grey lines.

## Summary

In this work, we demonstrated improvements of the depth of proteome coverage and quantification accuracy in the analysis of dia-PASEF data with an ion-mobility augmented version of DIA-NN (Demichev et al., 2020) and with spectral libraries generated using FragPipe (Kong et al., 2017; da Veiga Leprevost et al., 2020; Yu et al., 2020a), with both these workflow elements contributing to the better performance. We describe the identification of over 5200 proteins from low amounts (10ng) of a HeLa proteomic preparation, from the data acquired using a standard nanoLC gradient of 95 minutes. Further, when analysing runs acquired using 50ng of HeLa preparation, we identify over 700 extra proteins compared to the originally published results for 100ng injections. Moreover, when analysing the 4.8-minute Evosep One runs, we identify over 5000 proteins, and from just 200ng of HeLa preparation. In summary, by combining spectral library generation capabilities of FragPipe with TIMS-expanded version of DIA-NN, we provide an easy to use yet fast and powerful computational workflow that achieves high depth of coverage and excellent quantification precision in the analysis of dia-PASEF data.

**Figure S1.**
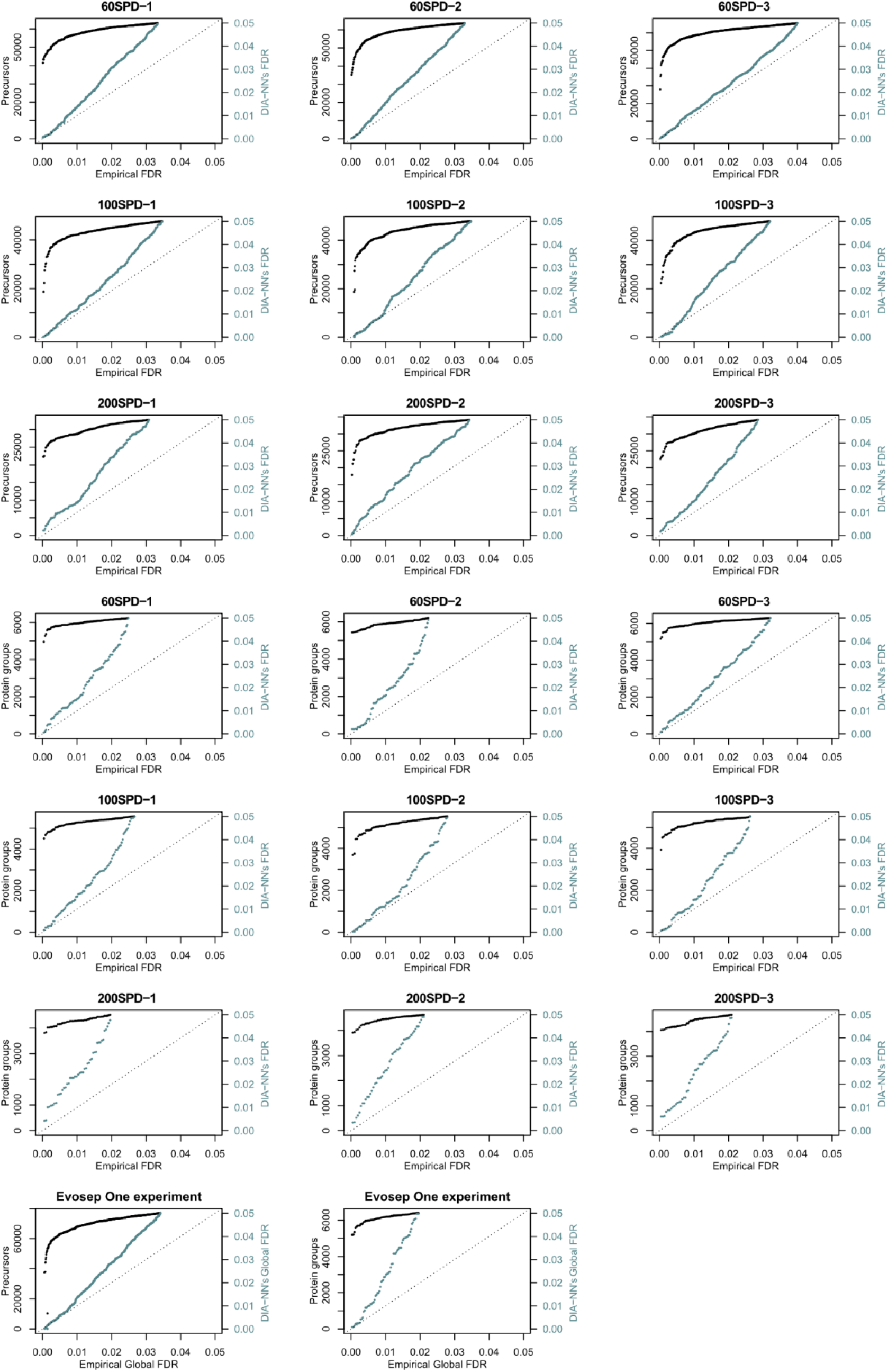
False-discovery rate validation. Evosep One acquisitions at different throughputs (60 SPD - 200 SPD) were analysed. Precursor and protein numbers (black) as well as FDR values reported by DIA-NN (grey) plotted against the empirical FDR estimates obtained (Methods) using a two-species human-*Arabidopsis* spectral library (Muntel et al., 2019), for individual runs (run-specific FDR) as well as for the entire experiment (Global FDR). Identifications were ordered by their q-values and visualisation performed for precursors starting with rank 10000 and proteins starting with rank 1000. Each point corresponds to an *Arabidopsis* precursor or protein called in a human sample. The benchmark reveals that FDR values reported by DIA-NN are highly conservative. Further, the benchmark demonstrates that DIA-NN can achieve < 0.1% false discovery rates, at both precursor and protein level, regardless of the data complexity (gradient length).

**Figure S2.**
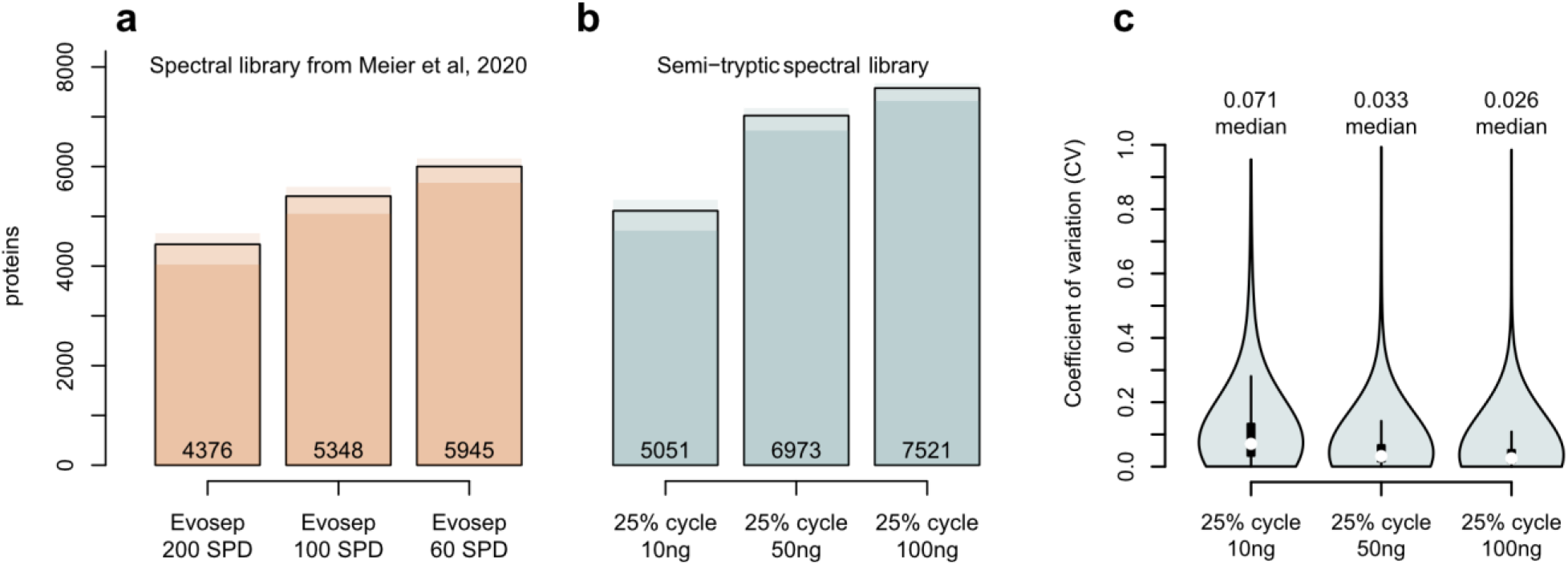
**a** Numbers of unique proteins, identified using the spectral library from the previous dia-PASEF workflow (Meier et al., 2020). **b** Numbers of protein groups identified in the 25% duty cycle runs with different injection amounts using a semi-tryptic spectral library and **c** the respective CV value distributions. Numbers of proteins identified in 1, 2 or all 3 replicates are shown with different color shades.

## Methods

### Spectral library generation in FragPipe

We used FragPipe computational platform (version 15) with MSFragger (Kong et al., 2017; Yu et al., 2020a) (version 3.2), Philosopher (da Veiga Leprevost et al., 2020) (version 3.4.13), and EasyPQP (https://github.com/grosenberger/easypqp; version 0.1.9) components to build spectral libraries. Peptide identification from tandem mass spectra (MS/MS) was done using MSFragger search engine, using either raw (.d) files (Evosep and nanoflow HeLa dataset) or MGF files (two-organism dataset) as input. Protein sequence databases *H. sapiens* (UP000005640) or *S. cerevisiae* (UP000002311) from UniProt (reviewed sequences only; downloaded on Feb. 15, 2021) and common contaminant proteins, containing in total 20421 (*H. sapiens*) and 6165 (*S. cerevisiae*) sequences were used. Reversed protein sequences were appended to the original databases as decoys. For the MSFragger analysis, both precursor and (initial) fragment mass tolerances were set to 20 ppm. Spectrum deisotoping (Teo et al., 2021), mass calibration, and parameter optimization (Yu et al., 2020b) were enabled. Enzyme specificity was set to ‘stricttrypsin’ (i.e. allowing cleavage before Proline), and either fully enzymatic or semi-enzymatic peptides were allowed (see Results for detail). Up to two missed trypsin cleavages were allowed. Isotope error was set to 0/1/2. Peptide length was set from 7 to 50, and peptide mass was set from 500 to 5000 Da. Oxidation of methionine, acetylation of protein N-termini, −18.0106 Da on N-terminal Glutamic acid, and −17.0265 Da on N-terminal Glutamine and Cysteine were set as variable modifications. Carbamidomethylation of Cysteine was set as a fixed modification. Maximum number of variable modifications per peptide was set to 3.

For each analysis, the MS/MS search results were further processed using the Philosopher toolkit (da Veiga Leprevost et al., 2020). First, MSFragger output files (in pepXML format) were processed using PeptideProphet (Keller et al., 2002) (with the high-mass accuracy binning and semi-parametric mixture modeling options) to compute the posterior probability of correct identification for each peptide to spectrum match (PSM). The resulting output files from PeptideProphet (also in pepXML format) were processed using ProteinProphet (Nesvizhskii et al., 2003) to assemble peptides into proteins (protein inference) and to create a combined file (in protXML format) of high confidence proteins groups and the corresponding peptides assigned to each group. The combined protXML file was further processed using Philosopher Filter module as follows. Each identified peptide was assigned either as a unique peptide to a particular protein (or protein group containing indistinguishable proteins) or assigned as a razor peptide to a single protein (protein group) that had the most peptide evidence. The protein groups assembled by ProteinProphet, with the probability of the best peptide used as a protein-level score (Nesvizhskii, 2010), were filtered to 1% protein-level FDR using the picked FDR strategy (Savitski et al., 2015), allowing unique and razor peptides. The final reports were then generated and filtered at each level (PSM, ion, peptide, and protein) using the 2D FDR approach (Bern and Kil, 2011) (1% protein FDR plus 1% PSM/ion/peptide-level FDR for each corresponding PSM.tsv, ion.tsv, and peptide.tsv files).

Finally, PSM.tsv files, filtered as described above, along with the spectral files (original MGF files, or uncalibrated MGF files created by MSFragger when raw .d files were used as input to MSFragger) were used as input to EasyPQP for generation of the consensus spectrum libraries. In doing so, a peptide’s retention times in each fraction were non-linearly aligned (lowess method) by EasyPQP to a common iRT scale using the extended HeLa iRT calibration peptide set (Bruderer et al., 2016). Peptide’s ion mobility values in each run in the dataset were aligned to that from one of the runs in the dataset automatically selected as a reference run. The library was additionally filtered to keep only peptides contained in the Philosopher-generated peptide.tsv report file, ensuring that the final spectral library was filtered to 1% protein and 1% peptide-level FDR.

### Spectral library processing in DIA-NN

For the two-proteome human-yeast benchmark, the HeLa library was filtered to only include peptides present in the *in silico* tryptic digest of the human database and exclude peptides present in the tryptic digest of the yeast database and vice versa, by generating the annotation of the library using the “Reannotate” function in DIA-NN and discarding peptides matched to one or more proteins of the other species.

### DIA-NN configuration and dia-PASEF data processing

DIA-NN 1.7.15 was used for the benchmarks and was operated with maximum mass accuracy tolerances set to 10ppm for both MS1 and MS2 spectra. The two-proteome human-yeast benchmark was analysed with match-between-runs (MBR) enabled, similarly to the previous dia-PASEF workflow (Meier et al., 2020). Protein inference in DIA-NN was disabled (except for one benchmark as noted below), to use the protein groups assembled at the spectral library building stage in FragPipe. When reporting protein numbers and quantities, the Protein.Group column in DIA-NN’s report was used to identify the protein group and PG.MaxLFQ was used to obtain the normalised quantity. PSM tables (PSM.tsv files generated by Philosopher) contain accession numbers of all mapped proteins for each identified peptide, and this information was used to identify proteotypic peptides. In the benchmark for the numbers of unique proteins with the spectral library from the original dia-PASEF publication (Supp. Figure S2), the “Genes” column was used to count unique proteins (as gene products identified and quantified using proteotypic peptides only). For this, proteotypic peptides were annotated as such using the “Reannotate” option in DIA-NN. Quantification mode was set to “Robust LC (high precision)”. All other settings were left default. Following previously published recommendations (Rosenberger et al., 2017), and similarly to the previous dia-PASEF workflow (Meier et al., 2020), DIA-NN’s output was filtered at precursor q-value < 1% and global protein q-value < 1%.

Of note, one of the two-proteome human-yeast benchmark files (200113_AUR_dia-PASEF_HY_200ng_15ng_90min_Slot1-5_1_1636.d) could not be read correctly due to data corruption, with all frames (i.e. dia-PASEF scans) from 55877 onwards being discarded, which might have resulted in the benchmark results being very slightly worse.

### FDR validation benchmark using a two-species human-Arabidopsis library

The library (Muntel et al., 2019) (repository PXD013658, file “HumanThalianaDDAOnly (two-species FDR test).xls”) was filtered to only include unmodified peptides or peptides with carbamidomethylated cysteines. DIA-NN was instructed to replace all spectra and retention times in this library with in silico predicted ones, as well as generate in silico reference ion mobility (1/K0) values. This was done to eliminate any potential bias in spectral quality between human and plant peptides in this experimental library. “Precursor FDR” filter was set to 10% in DIA-NN settings.

The empirical FDR was then determined experimentally for each of the Evosep One runs with different gradient lengths, by counting *Arabidopsis* precursor/protein calls. Specifically, the FDR was determined as

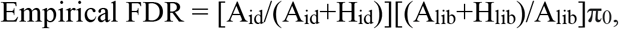

where A_id_ is the number of calls of *Arabidopsis* precursors or proteins at a particular score threshold, H_id_ - calls of human precursors or proteins, and A_lib_, H_lib_ - the respective numbers of precursors or proteins in the library. The π_0_ (‘prior probability of incorrect identification’, also known as PIT - <u>P</u>ercentage of <u>I</u>ncorrect <u>T</u>argets) correction factor (Käll et al., 2008; Storey, 2002) was calculated as

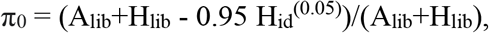

where H_id_^(0.05)^ is the number of human precursor or protein calls at q-value = 0.05. The numbers of precursors/proteins were obtained based on filtering the library for precursors within the mass range 400.0 - 1000.0 m/z (the range sampled in the Evosep One dia-PASEF runs under consideration). The empirical FDR was then plotted (Supp. Figure S1) against the precursor/protein FDR estimates reported by DIA-NN.

## Data availability

All the spectral libraries, PSM tables and DIA-NN’s reports, logs and the pipeline configuration file were deposited to an OSF (Open Science Framework) repository https://osf.io/3k2zx/?view_only=1750e62e939641ada2e6f2f81aa9e99d.

DIA-NN is freely available for download from https://github.com/vdemichev/DiaNN. FragPipe is freely available for download from https://fragpipe.nesvilab.org/. MSFragger is freely available under academic license and can be downloaded at http://msfragger.nesvilab.org/ or directly from FragPipe.

## Acknowledgements

We thank Sven Brehmer and Nagarjuna Nagaraj (Bruker) for helpful discussions and their advice on the analysis of dia-PASEF data. We thank Dmitry Avtonomov for his help with FragPipe development. This work was funded in part by NIH grants R01-GM-094231 and U24-CA210967 (to AIN), the BBSRC (BB/N015215/1 to MR, BB/N015282/1 to KSL), the Francis Crick Institute, which receives its core funding from Cancer Research UK (FC001134), the UK Medical Research Council (FC001134), and the Wellcome Trust (FC001134 and IA 200829/Z/16/Z to M.R), and the Ministry of Education and Research (BMBF), as part of the National Research Node ‘Mass spectrometry in Systems Medicine (MSCoresys), under grant agreement 031L0220A (MR).

